# Describing small angle scattering profiles by a limited set of intensities

**DOI:** 10.1101/2021.05.24.445439

**Authors:** Thomas D. Grant

## Abstract

Small angle scattering (SAS) probes the size and shape of particles at low resolution through the analysis of the scattering of X-rays or neutrons passing through a solution of particles. One approach to extracting structural information from SAS data is the indirect Fourier transform (IFT). The IFT approach parameterizes the real space pair distribution function (*P*(*r*)) of a particle using a set of basis functions, which simultaneously determines the scattering profile (*I*(*q*)) using corresponding reciprocal space basis functions. This article presents an extension of an IFT algorithm proposed by Moore which used a trigonometric series to describe the basis functions, where the real space and reciprocal space basis functions are Fourier mates. An equation is presented relating the Moore coefficients to the intensities of the SAS profile at specific positions as well as a series of new equations that describe the size and shape parameters of a particle from this distinct set of intensity values. An analytical real space regularizer is derived to smooth the *P*(*r*) curve and ameliorate systematic deviations caused by series termination, commonly used in IFT methods though not described in Moore’s original approach, which is particularly susceptible to such effects. The algorithm is provided as a script, *denss.fit_data.py*, as part of the DENSS software package for SAS, which includes both command line and interactive graphical interfaces. Results of the program using experimental data show that it is as accurate, and often more accurate, than existing tools.

## 1. Introduction and Overview

Small angle scattering (SAS) yields structural information at low resolution about the size and shape of particles in solution. X-rays or neutrons scattering from freely tumbling particles in solution exhibit rotational averaging in reciprocal space, resulting in isotropic scattering profiles collected on 2D detectors. This rotational averaging results in the loss of information describing the three-dimensional structure of the particle. The scattering of a molecule, *I*(*q*), where *q* is the momentum transfer (defined in Appendix A), is determined by its three-dimensional scattering length density function, thus SAS profiles can be calculated directly from known atomic structures. However, due to the spherical averaging of the intensities, the inverse problem of calculating a unique three-dimensional structure from SAS profiles is not possible. Nonetheless, structural information describing global properties of size and shape can be obtained through analysis of the SAS profile.

While unique three-dimensional real space information cannot be obtained directly from a SAS profile, a Fourier transform of the reciprocal space intensity profile yields the set of pair distances in the particle, known as the pair distribution function, or *P*(*r*). However, due to limitations caused by termination of higher order scattering data to a finite *q* range, uncertainties in intensity measurements, and systematic errors, direct calculation of the Fourier transform yields *P*(*r*) functions with large systematic deviations (Glatter, 1977; Moore, 1980; Hansen & Pedersen, 1991; Svergun, 1992; Svergun & Pedersen, 1994). One popular approach to extracting this structural information from SAS profiles is the indirect Fourier transform (IFT) proposed by Glatter (Glatter, 1977). In this approach, a set of basis functions is used to parameterize the *P*(*r*) function. The weights of these basis functions are then adjusted to optimize the fit of the corresponding intensity function to the experimental scattering profile.

One such IFT algorithm proposed by Moore (Moore, 1980) leverages information theory (Shannon, 1948) to describe a set of basis functions defined by the maximum particle dimension, *D.* Moore uses a trigonometric series to define a function *Q*(*r*) = *P*(*r*)/*r*. This definition resulted in a convenient relationship between the real space Q(r) and the reciprocal space *U*(*q*) = *qI*(*q*), where the two are Fourier mates. Key to Moore’s approach (and other IFT methods (Glatter, 1977; Svergun, 1992)) is that the coefficients of the series terms define both the real space and reciprocal space profiles, using the appropriate basis functions. Least squares can be used to determine the coefficients and the associated standard errors by minimizing the fit to the experimental scattering profile (Appendix A). This approach has the advantage of providing the necessary information on the variances and covariances of the coefficients to determine the errors on each coefficient. Moore showed using Shannon information theory that the number of coefficients that can be determined from the data is the number of independent pieces of information that the data are able to describe about the particle. Moore derived a series of equations relating the coefficients to commonly used SAS parameters such as the forward scattering intensity, *I*(0), the radius of gyration, *R_g_*, and the average vector length, 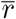, along with error estimation for each parameter. One advantage of Moore’s approach over others is that a separate regularizing function is not explicitly required to smooth the *P*(*r*) curve due to the use of the sine series (Moore, 1980). However, in practice with experimental data, it has been found that Moore’s approach is often more susceptible to large oscillations in the *P*(*r*) curve due to series termination (Svergun & Pedersen, 1994; Hansen & Pedersen, 1991), likely due to the lack of a regularizing function. Such regularizing functions have been shown to be effective at smoothing the *P*(*r*) curves calculated using Moore’s approach (Tully *et al*., 2021; Rambo, 2021).

Here we extend Moore’s derivation to relate the Moore coefficents to specific intensity values such that each term in the series is now weighted by a corresponding intensity, termed *I_n_* (Appendix A). We present equations for calculating a variety of commonly used SAS parameters and their associated errors from the *I_n_*’s. Additionally, we derive a modified equation for least squares minimization taking into account an analytical regularization of the *P*(*r*) curve. We provide open-source software with convenient interfaces for performing all of the presented calculations, including a novel approach to estimating parameters sensitive to systematic errors. Finally, we describe the results using both simulated and real experimental data and compare with current state-of-the-art software tools.

## 2. Theoretical Background

### 2.1. Extension of Moore’s IFT

Moore’s use of Shannon information theory to define *I*(*q*) resulted in a selection of *q* values, namely *q_n_* = *nπ*/*D*, termed “Shannon channels” (Feigin & Svergun, 1987; Svergun & Koch, 2003; Rambo & Tainer, 2013). The intensities at *q_n_*, i.e. *I_n_* = *I*(*q_n_*), therefore become important values as they determine the Moore coefficients, *a_n_*, and thus similarly can be used to completely describe the low-resolution size and shape of a particle obtainable by SAS. In Appendix A we derive the mathematical relationship between *I_n_* and *a_n_* which results in the following general equation for *I*(*q*) as a function of the intensity values at the Shannon points:

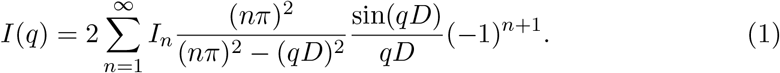

Defining basis functions *B_n_* as

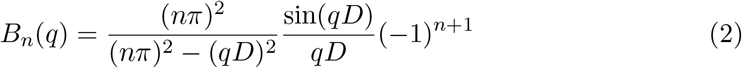

*I*(*q*) can now be expressed as a sum of the basis functions *B_n_* weighted by the intensity values at *q_n_*

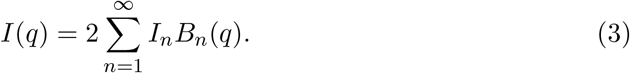

As in Moore’s original approach, the *B_n_* functions are determined by the maximum dimension of the particle, *D. B_n_*’s for *D* = 50 Å are illustrated in Figure 1. The *P*(*r*) function can be represented using the series of *I_n_* values as

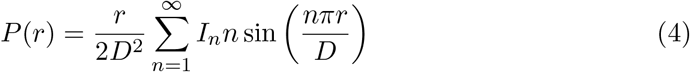

**Fig. 1.**
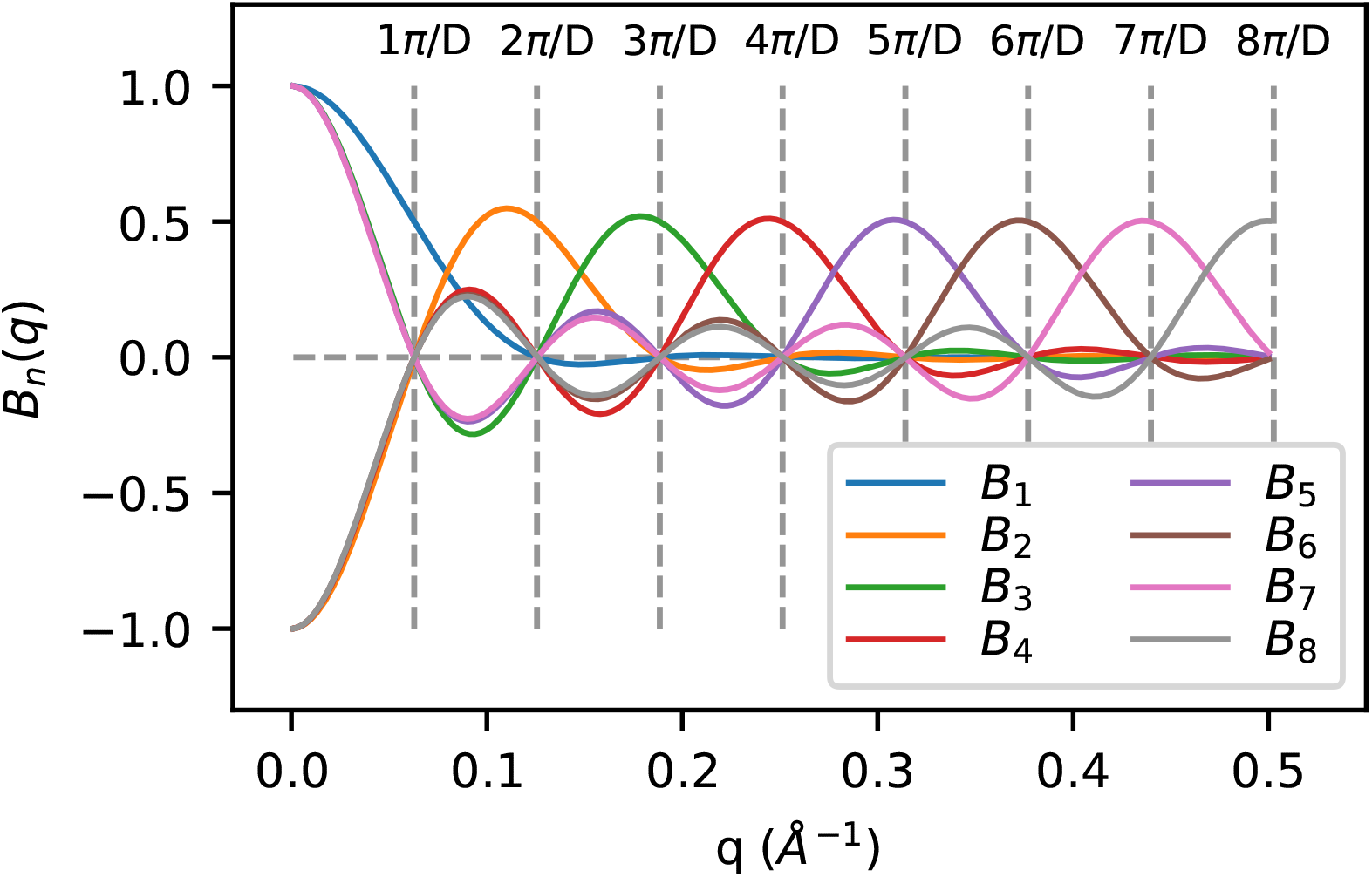
Plot of reciprocal space basis functions, *B_n_*, for any particle of size *D* = 50 Å. Vertical dashed lines show the locations of the Shannon points, *q_n_*.

(Appendix A) or by defining real space basis functions *S_n_* as follows:

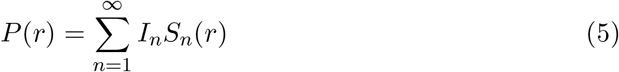

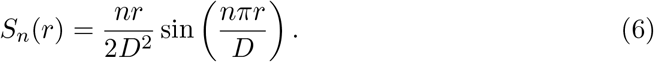

Least squares can be used to determine optimal values for each *I_n_* from the oversampled experimental SAS profile along with error estimates for each taking into account the variances and covariances of the coefficients. These terms can then be used to calculate the corresponding *I*(*q*) and *P*(*r*) curves using equations 1 and 4 as well as the associated errors (Appendix A).

The maximum particle dimension, *D*, is required for determining the *q_n_* values associated with the *I_n_*’s. Estimates for the true value of D that are too small will result in *B_n_*’s that lack sufficiently high frequencies to adequately reconstruct *I*(*q*). Estimates of *D* that are too large will result in overfitting the data. Moore found that testing increasing values of *D* yielded improved fits to the experimental *I*(*q*) function and used *χ*^2^ (Appendix A) to estimate the true value of *D* by selecting the smallest *D* value that minimizes *χ*^2^ while avoiding larger *D* values that result in overfitting (Moore, 1980). An alternative method is to estimate *D* from the *P*(*r*) curve by first guessing a reasonable value for *D*, such as 3.5*R_g_* or larger, fit *I*(*q*) and calculate the *P*(*r*) curve, and then estimate the true value of *D* based on where *P*(*r*) gradually falls to zero.

### 2.2. Derivation of Parameters from I_n_’s

Similar to what Moore described for the an coefficients, since the *I_n_*’s contain all the information present in *I*(*q*), quantities that can be derived from *I*(*q*) can also be derived directly from the *I_n_*’s. For example, to determine the forward scattering intensity, *I*(0), we take the limit of equation 1 as *q* approaches zero to yield

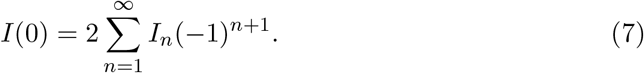

Equation 7 demonstrates a simple relationship between the forward scattering of a particle and the *I_n_*’s. Note that the particle dimension, *D*, is not explicitly present in equation 7. Figure 2 illustrates the relationship of the *I_n_*’s and *I*(0).

**Fig. 2.**
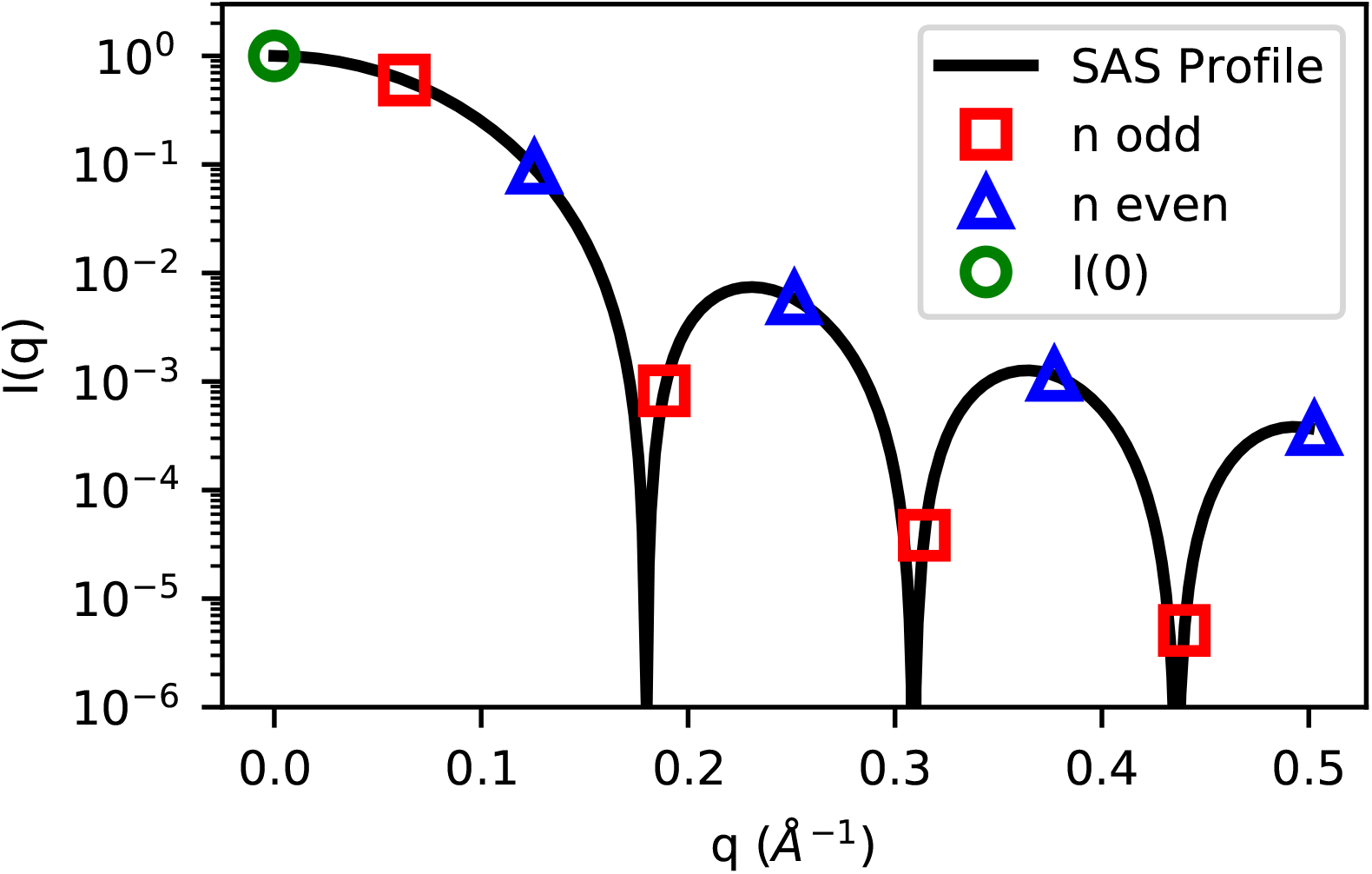
Plot of example scattering profile showing the relationship of *I_n_*’s and *I*(0). Odd *I_n_*’s are shown as red squares, while even *I_n_*’s (which have a multiplication factor of −1 in equation 7) are shown as blue triangles. Equation 7 says that twice the total sum of the red squares and (negative) blue triangles is equal to the forward scattering, *I*(0), shown as a green circle.

The forward scattering of a particle is not directly measured in an experiment due to its coincidence with the incident beam and is thus typically estimated as an extrapolated value from low *q* data points or by integration of the *P*(*r*) function. Equation 7 provides an alternative method to measuring the forward scattering of a particle directly from the data through the sum of the *I_n_*’s. While equation 7 is defined as a sum from *n* = 1 to infinity, typical experimental setups only provide data for the first 10 to 30 Shannon channels, depending on the size of the particle. Thus in practice equation 7 yields an estimate of the forward scattering rather than an exact measurement. However, since the vast majority of the scattering intensity present in the profile occurs within these 10 to 30 Shannon channels, equation 7 should provide an accurate estimate of the forward scattering for most particles and experimental setups.

Other parameters can similarly be derived (Appendix B). For example, *R_g_* can be estimated from the *I_n_*’s as

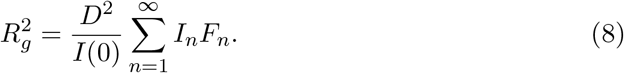

where

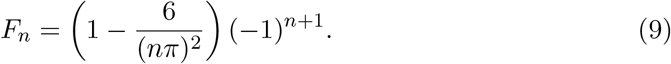

Another parameter describing particle size is the average vector length in the particle, 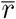, which can be estimated from the *I_n_*’s as

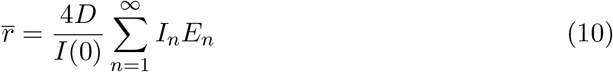

where

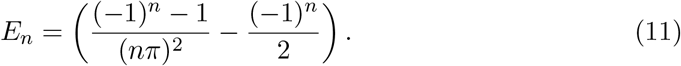

The Porod invariant, *Q*, is defined as the integrated area under the Kratky plot (Porod, 1982) which can be described in terms of the *I_n_*’s as

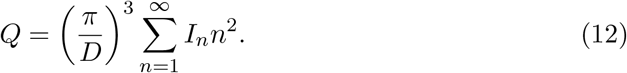

The Porod volume can then be calculated using the Porod invariant (Appendix B) (Porod, 1982). The Porod volume is commonly used to estimate molecular weight for globular biological macromolecules. More recently Rambo and Tainer (Rambo & Tainer, 2013) derived a new SAS invariant termed the Volume of Correlation, *V_c_*, whose unit is *length*^2^ and is related to the correlation length of the particle, *ℓ_c_*. *V_c_* can be used to estimate molecular weight from macromolecules that may be either globular or flexible (Rambo & Tainer, 2013). *V_c_* can be estimated from the *I_n_*’s as

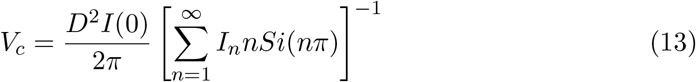

where *Si*(*nπ*) is the Sine Integral. The length of correlation can similarly be calculated as

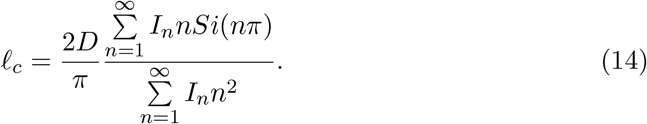

Since the variances and covariances of the *I_n_*’s are known from the least squares minimization, error propagation can be used to determine the associated uncertainties for each of the parameters described above (Appendix B).

### 2.3. Regularization of P (r)

The original IFT proposed by Glatter (Glatter, 1977) as well as other IFTs (Svergun, 1992; Vestergaard & Hansen, 2006), makes use of regularization of the *P*(*r*) curve similar to the general method of Tikhonov regularization (Tikhonov & Arsenin, 1977). The goal is to use the knowledge that *P*(*r*) functions for most particle shapes are smooth to generate curves that are free of strong oscillations from series termination and are relatively stable to statistical errors. Rather than minimize *χ*^2^ directly, a new function, *T*, is minimized taking into account the smoothness of the *P*(*r*) curve according to equation 15:

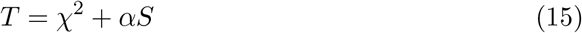

where *S* is the regularizing function which can take different forms, and *α* is a Lagrange multiplier that acts as a weight to determine the strength of the smoothing. Larger *α* leads to a smoother *P*(*r*) function, but may result in a worse fit of *I*(*q*) to the experimental data. The IFT method by Moore has been shown to be more susceptible than other IFT methods to oscillations in the *P*(*r*) curve (Hansen & Pedersen, 1991; Svergun & Pedersen, 1994), most likely due to the lack of a regularizing function. We provide a detailed derivation of an analytical regularization of *P*(*r*) using *I_n_* values in Appendix C.

As for other similar IFT methods utilizing regularization, a suitable choice of *α* must be found to optimize the smoothness of the *P*(*r*) curve and the fit to the experimental data. Various methods for selecting the optimal value for *α* have been proposed, including via point of inflection (Glatter, 1977), Bayesian methods (Vestergaard & Hansen, 2006) and using perceptual criteria (Svergun, 1992). We describe our approach in section 2.4 below.

Equation 3 assumes a sum from *n* = 1 to infinity. However, data are only collected to the maximum *q* value allowed by experiment, *q_max_*. The lack of data for *q* > *q_max_* implicitly corresponds to setting the *I_n_* values to zero for those data points where *n* > *n_max_* (where *n_max_* = int(*q_max_D/π*), i.e. the largest index in the series). The regularization often results in poorer fits of the intensity profile at higher experimental *q* values with increasing *α* due to this implicit bias of *I_n_* values for *n* > *n_max_* towards zero. In order to remove this bias and allow for the *I_n_* values at *n* > *n_max_* to be unrestrained, *I_n_* values for *n* > *n_max_* are allowed to float (calculated up to 3*n_max_*). It is important to note that the number of Shannon channels that can be reliably extracted from the data are dictated largely by the quality of the data in addition to the *q* range, as described in (Konarev & Svergun, 2015).

### 2.4. Implementation

Tools for performing the least squares fitting of *I_n_*’s to experimental data, calculation of parameters and errors, and regularization of *P*(*r*), have been developed using Python, NumPy, and SciPy (Harris *et al*., 2020; Virtanen *et al*., 2020) and are provided open source through the DENSS suite of SAS tools (Grant, 2018) (https://github.com/tdgrant1/denss). The primary interface to use this algorithm is the *denss.fit_data.py* Python script. To enable ease of use, in addition to the command line interface, an interactive graphical user interface (GUI) has been developed using the Matplotlib package (Figure 3) (Hunter, 2007).

**Fig. 3.**
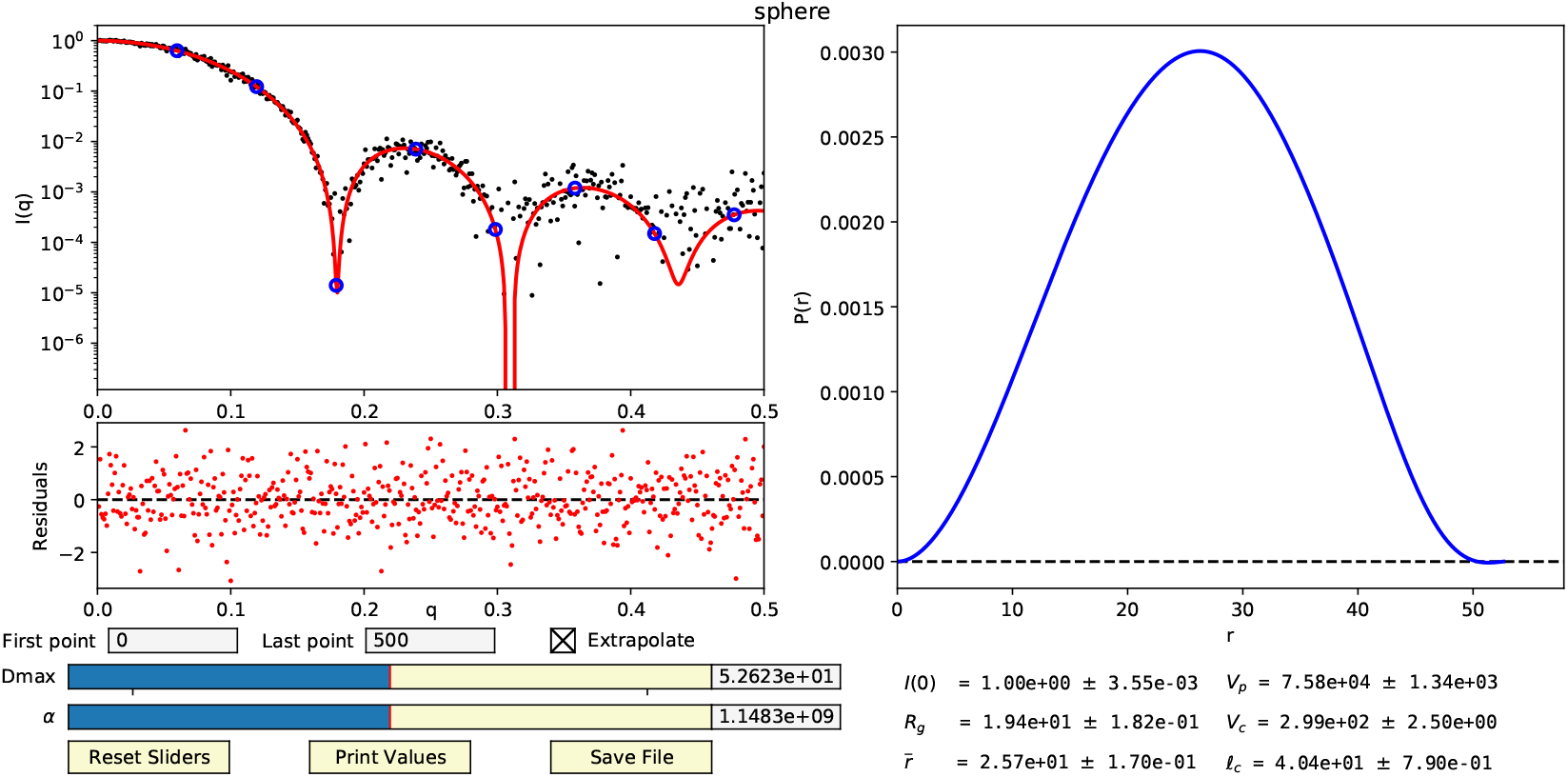
Interactive GUI display from the *denss.fit_data.py* script. The upper left panel shows the experimental *I*(*q*) curve as black circles, fitted *I_n_* values as blue circles, and the fitted *I_c_*(*q*) calculated from the *I_n_*’s as a red curve on a semilog plot. The residuals of the experimental and calculated intensity curves are shown below the intensity plot. The panel on the right shows the *P*(*r*) curve calculated from the *I_n_*’s. Input text boxes on the bottom left allow for trimming data points at the beginning or end of the curve along with a checkbox to disable the calculation of intensities at high *q* values. Interactive sliders for *D_max_* and *α* are also displayed along with corresponding input boxes for manual entry. The bottom right of the window shows size parameters calculated from the *I_n_*’s and associated uncertainties. Buttons for resetting the sliders, printing the size parameters, and saving the results are shown on the bottom left.

#### 2.4.1. Automatic estimation of D

To assist users, upon initialization of the script, the experimental data are loaded and estimates of *D* and *α* are automatically calculated. To automatically estimate *D*, an initial estimate of *D* is calculated that is likely to be significantly larger than the actual *D*. This subsequently enables a more accurate estimation of *D* where *P*(*r*) falls to zero. An initial value of *D* = 7*R_g_* is used as this should ensure a large enough value given a variety of particle shapes (Petoukhov *et al*., 2007; Grant *et al*., 2015). An initial rough estimate of *R_g_* is first calculated using Guinier’s equation (Guinier *et al*., 1955) with the first twenty data points. In cases where that estimate fails (e.g. due to excessive noise or a positive slope of the Guinier plot), the Guinier peak method is instead used (Putnam, 2016). The *I_n_*’s are then calculated from the experimental data using the regularized least squares approach outlined in Appendix C, setting *α* = 0 to optimize the fit to the data. After the initial *I_n_*’s are calculated, the corresponding *P*(*r*) function often suffers from severe ripples caused by Fourier termination effects due to the finite range of data, as described above, making it difficult to estimate *D* where *P*(*r*) falls to zero. To alleviate this effect, a Hann filter, which is a type of Fourier filter (Blackman & Tukey, 1958), is applied to remove the Fourier truncation ripples from *P*(*r*). *D* is then calculated from this filtered *P*(*r*) curve as the first position, *r*, where *P*(*r*) falls below 0.01*P_max_* after the maximum, where *P_max_* is the maximum value of the filtered *P*(*r*). This new *D* value is then used to recalculate the *I_n_*’s for the best fit to the experimental scattering profile. In addition to automatically estimating *D* directly from the data, users can manually enter an initial estimate of *D* to begin.

#### 2.4.2. Automatic estimation of α

Next, the optimal *α* is estimated which yields *I_n_* values corresponding to a smooth *P*(*r*) function while still resulting in a calculated *I*(*q*) curve that fits the experimental data. First, the best *χ*^2^ value possible is calculated by setting *α* = 0 and using the *D* value estimated in the previous step. Then, various values of *α* are scanned, from 10^-20^ to 10^20^ in logarithmic steps of 10^1^. This wide range is used to accommodate a variety of different scattering profiles covering a range of signal to noise values. At each step the *χ*^2^ is calculated. The optimal *α* is chosen by interpolating where 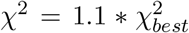, i.e., where the *χ*^2^ rises to 10% above the best possible value.

#### 2.4.3. Interface

The GUI mode of the script displays a plot of the intensities on a semilog y-axis and plots the experimental data, *I_e_*(*q*), and the initial fit, *I_c_*(*q*), calculated from the *I_n_*’s at the experimental *q* (Figure 3). The script additionally calculates *I_c_*(*q*) at *q* values extrapolated to *q* = 0. Users can alternatively provide a set of desired *q* values to calculate *I_c_*(*q*) as an ASCII text file when starting the program. The residuals, (*I_e_*(*q_i_*) — *I_c_*(*q_i_*))/*σ_i_*, are also displayed to assist in assessing the quality of the fit. Next to the plot of intensities, the *P*(*r*) curve calculated from the *I_n_*’s is also displayed. In addition to manually entering new *D* and *α* values in input text boxes in the GUI, interactive sliders are available to change the *D* and *α* values, which automatically update the plots as they are adjusted. Users can also change the beginning and ending data points if desired to remove outlier data points that often occur at either end of the experimental profile or disable the calculation of intensities for *q* > *q_max_*. Several of the parameters described above, including 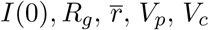, and *ℓ_c_*, along with associated uncertainties, are calculated from the *I_n_*’s and displayed on the GUI. These parameters are updated interactively whenever *D* or *α* are changed.

#### 2.4.4. Calculation of V_p_, V_c_, and ℓ_c_

Particular care must be taken when estimating parameters that are sensitive to systematic errors in high *q* data points, such as *V_p_, V_c_*, and *ℓ_c_*. In practice, direct estimation of these parameters using the equations described above may yield unstable results, even with regularization. Porod’s law is based on the assumption that all scattering comes from the surface of a particle, resulting in an asymptotic intensity decay proportional to *q*^-4^ (Porod, 1982), giving rise to the ability to estimate values such as the Porod volume, *V_p_*. In practice, shape scattering contributes significantly (Rambo & Tainer, 2011), as do systematic errors caused by inaccurate background subtraction (Manalastas-Cantos *et al*., 2021), resulting in poor estimation of these parameters without correction. To deal with this, many algorithms impose an artificial constant subtraction to force the Porod decay which have proven effective at providing accurate estimates of particle volume (Manalastas-Cantos *et al.*, 2021). However, different algorithms have different methods for calculating the constant to subtract and for determining the fitting region where these calculations are performed, and there is often subjectivity involved in selecting the appropriate “Porod region” (Rambo & Tainer, 2011; Neto *et al*., 2021). To avoid such issues with constant subtraction altogether, we have developed a different approach. In our approach, we take advantage of the regularization provided above by intentionally *over* smoothing using a large *α*. Oversmoothing has the effect of removing shape scattering while simultaneously enforcing a decay similar to Porod’s law of *q*^-4^, making the resulting scattering profile more consistent with the assumptions of the Porod law. To do this, we multiply *α* by a factor of 10, which in our tests with experimental data resulted in the most accurate and robust results (see Results section below). We also limit the *q* range to 8/*R_g_*, which has previously been shown to be a reasonable cutoff for calculating Porod volume (Manalastas-Cantos *et al*., 2021; Neto *et al*., 2021). Note that this oversmoothing is only applied for calculation of the three parameters mentioned above and their associated errors and does not affect the actual fit of the scattering profile, *P*(*r*) curve, or other parameters.

#### 2.4.5. Output

Finally, upon exiting the script, the experimental data and calculated fit of the intensities are saved in a file with the calculated parameter values saved in the header. The corresponding *P*(*r*) curve is also saved. In addition to providing the *denss.fit_data.py* script as an interface to the algorithm described above, other scripts in the DENSS package also utilize this algorithm, including denss.py and denss.αll.py, to allow automatic fitting of the data and estimation of *D* and *α* when using these programs for *ab initio* 3D density reconstructions.

## 3. Results

One of the few shapes for which an analytical scattering equation has been derived is the solid sphere (Rayleigh, 1911; Porod, 1982). Since the equation of scattering for a sphere is known exactly, the *I_n_*’s for a sphere can be calculated directly (Appendix D), resulting in equation 16

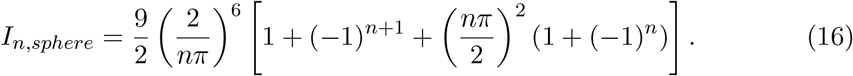

Note that the radius of the sphere, *R*, does not enter into equation 16. Interestingly, the odd *I_n_*’s for a sphere decay exactly as *q*^-6^ and the even *I_n_*’s decay exactly as *q*^-4^. The decay of intensity at higher angles proportional to *q*^-4^ is described by Porod’s Law (Porod, 1982) above, generally an approximation for most globular particles but here derived analytically for a sphere for even *I_n_*’s. All parameters outlined above, including *R_g_*, volume, etc., can be calculated analytically using equation 16, resulting in well-known equations for solid spheres (Appendix D). In Figure 4 the scattering profile for a sphere of radius 25 Å with added Gaussian noise (*I_e_*(*q*)) is shown with the fitted In values and the recovered *I_c_*(*q*) profile. Eight Shannon points were used to fit the data, from which size parameters were calculated using the fitted In values, shown in Table 1. The *I_n_* values can also be used to calculate the *P*(*r*) curve, *P_c_*(*r*), shown in Figure 5 along with the exact *P*(*r*) curve for a sphere (Porod, 1982) (Appendix D).

**Fig. 4.**
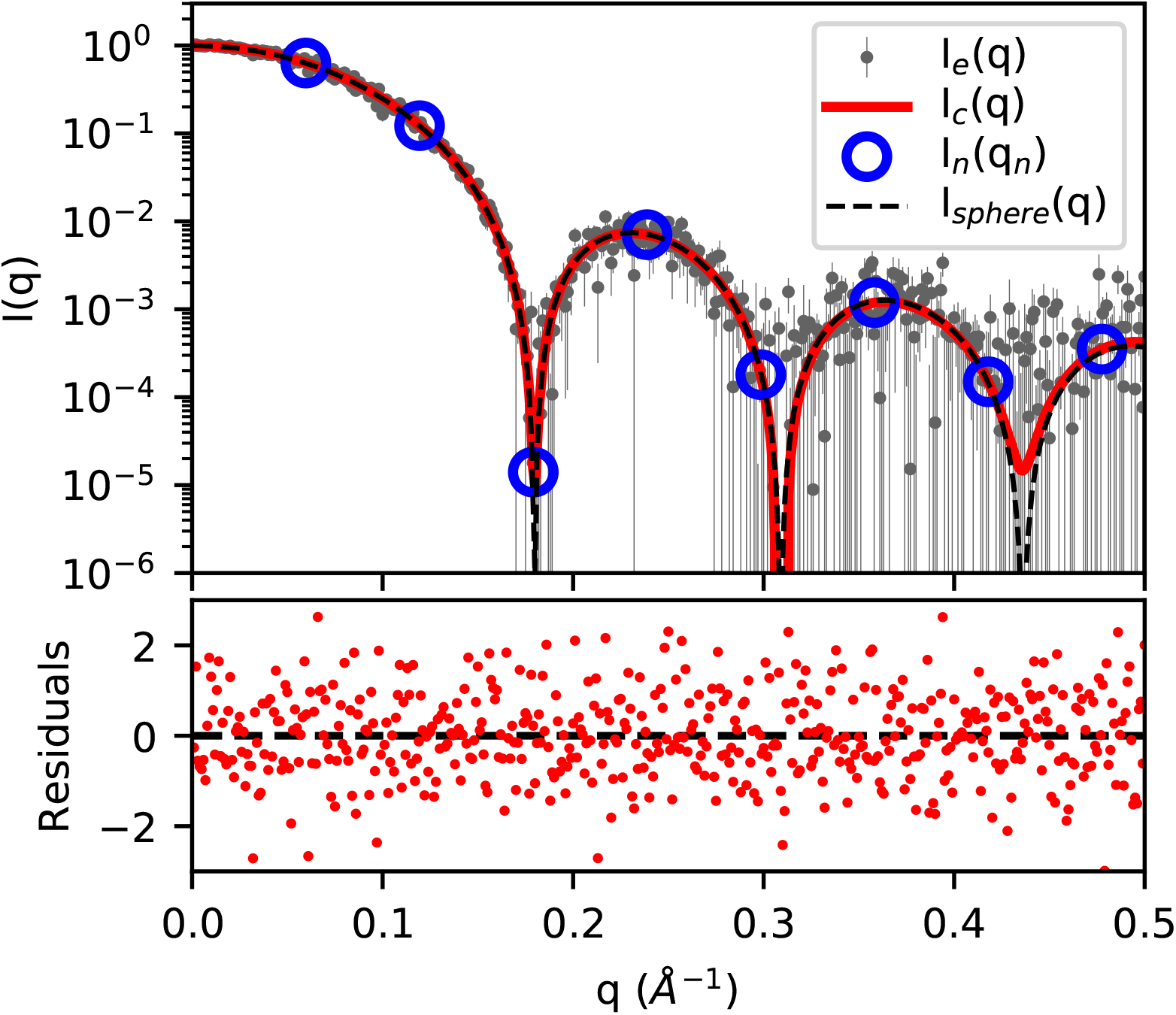
Plot of calculated intensity curve, *I_c_*(*q*) (red line), fitted to simulated noisy intensity values, *I_e_*(*q*) (gray dots with error bars), for a sphere of radius 25 A. The blue circles show the Shannon intensities, *I_n_*(*q_n_*), and the black dashed line shows the exact scattering profile of the sphere, *I_sphere_*(*q*). The bottom plot shows the residuals of the experimental data to the calculated profile.

**Fig. 5.**
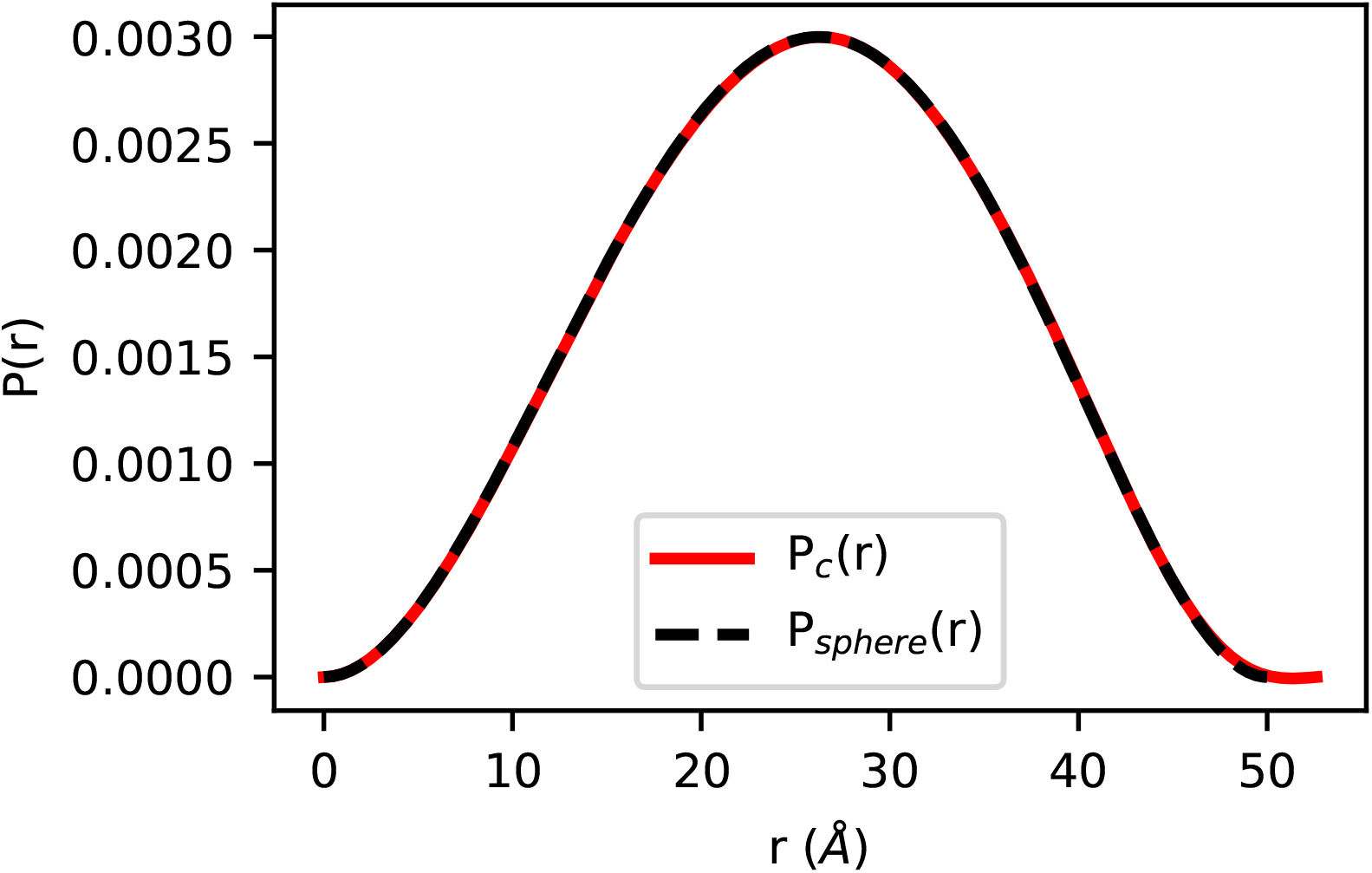
Plot of calculated *P_c_*(*r*) curve from *I_n_* values fitted to simulated noisy intensity values for a sphere of radius 25 Å. The exact *P*(*r*) curve for the sphere, *P_sphere_*(*r*) is also plotted as a dashed line.

**Table 1.**
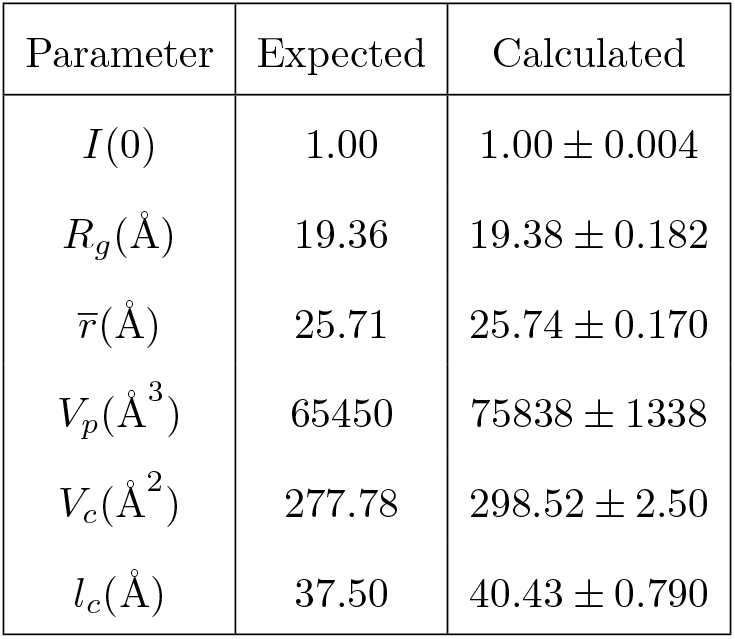
Table of parameters calculated from I_n_ values for the sphere profile shown in Figure 4. The columns correspond to expected parameter values using an infinite number of Shannon channels, and the recovered values calculated from the fit.

| Parameter | Expected | Calculated |
| --- | --- | --- |
| $I(0)$ | 1.00 | $1.00 \pm 0.004$ |
| $R_g(\text{\AA})$ | 19.36 | $19.38 \pm 0.182$ |
| $\bar{r}(\text{\AA})$ | 25.71 | $25.74 \pm 0.170$ |
| $V_p(\text{\AA}^3)$ | 65450 | $75838 \pm 1338$ |
| $V_c(\text{\AA}^2)$ | 277.78 | $298.52 \pm 2.50$ |
| $l_c(\text{\AA})$ | 37.50 | $40.43 \pm 0.790$ |

Data from publicly accessible databases for experimental SAS data, such as BIOISIS (https://www.bioisis.net) and SASBDB (Valentini *et al*., 2014), are particularly useful for verification and testing of algorithms such as described here. To test *denss.fit_data:py*. on experimental data sets, we downloaded two datasets from the benchmark section of the SASBDB online database, in particular SASDFN8 (apoferritin) and SASDFQ8 (bovine serum albumin) (Graewert *et al*., 2020). Automated estimates of *D* and *α* were suitable for accurate fitting and parameter estimation as indicated by the plot of residuals and comparison with the published parameter values (Figure 6). Best fits are achieved when setting *α* = 0, as expected, and increasing *α* results in smoother *P*(*r*) plots. High quality fits and smooth *P*(*r*) curves can be obtained simultaneously with an appropriate *α* (Figure 6), while setting *α* to too large of a value results in poorer fits to the intensity profile. Similar to other IFT methods, a balance must be struck to select the optimal *α* value resulting in the smoothest *P*(*r*) function possible while still enabling good quality fit of *I*(*q*).

**Fig. 6.**
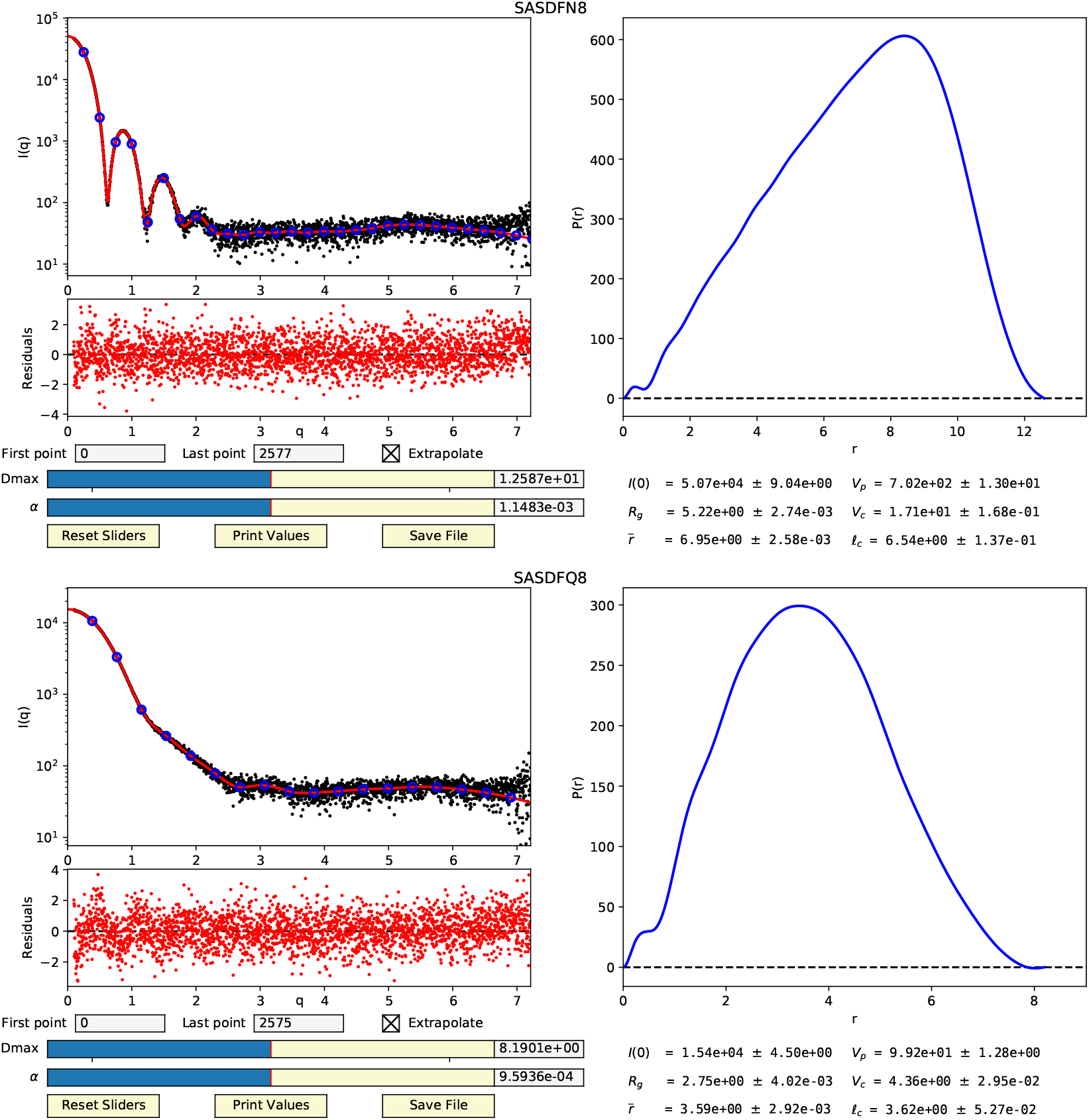
Fitting of *I_n_* values to real experimental datasets while using regularization results in good quality fits, smooth *P*(*r*) curves, and accurate parameter estimation. GUI displays of SASDFN8 (top) and SASDFQ8 (bottom). Note that the experimental *q* values were given in nanometers, resulting in nanometer units for parameters displayed.

To compare the parameter estimates with other software, we used DATGNOM from the ATSAS 3.0 package to estimate *R_g_* and *I*(0), DATPOROD to estimate *V_p_*, and DATVC to estimate *V_c_* from these two datasets (Manalastas-Cantos *et al*., 2021). A comparison of parameter values calculated by DATGNOM/DATPOROD/DATVC and *denss:fit_data:py* are shown in Table 2. Overall, and very importantly for community standards, the values are similar between the two different methods (~0.1% difference for *R_g_* and *I*(0) and ~3% difference for *V_p_* and *V_c_*). To verify that the error bounds are estimated correctly, we followed the protocol outlined in (Manalastas-Cantos *et al*., 2021) to use the DATRESAMPLE program to generate 1000 resampled scattering profiles from the two SASBDB datasets. This allows the calculation of parameters from each resampled profile and subsequently an estimate of the statistical errors based on the standard deviation of the parameter values, for comparison with the errors estimated by the programs. The results of this analysis are also shown in Table 2. The analysis shows that *denss.fit_data.py* produces similar or smaller statistical errors compared to estimated errors, suggesting the estimated errors should be considered an upper bound and the statistical errors likely less, whereas the statistical errors appear to be underestimated by DATGNOM (note, only *R_g_* and *I*(0) have estimated errors reported). It is noteworthy that the statistical errors on *R_g_* and *I*(0) are smaller from *denss.fit_data.py* (2 to 5-fold smaller) than from DATGNOM, while the statistical errors on *V_p_* and *V_c_* are about 2-fold smaller from DATG- NOM/DATPOROD/DATVC.

**Table 2.** Comparison of parameter values calculated from experimental datasets SASDFN8 (DFN8) and SASDFQ8 (DFQ8) using DATGNOM/DATPOROD/DATVC or denss.fit_data.py. Columns correspond to the value calculated for each parameter (Value) and either the estimated (± (Est.)) or statistical (± (Stat.)) errors as described in the text.

| Dataset | Parameter | DATGNOM/DATPOROD/DATVC |  |  | denss.fit_data.py |  |  |
| --- | --- | --- | --- | --- | --- | --- | --- |
| | | Value | ± (Est.) | ± (Stat.) | Value | ± (Est.) | ± (Stat.) |
| DFN8 | $R_g(nm)$ | 5.216e+00 | 5.451e-04 | 2.931e-03 | 5.222e+00 | 2.835e-03 | 5.933e-04 |
| DFN8 | $I(0)$ | 5.063e+04 | 9.836e+00 | 2.306e+01 | 5.070e+04 | 9.173e+00 | 9.061e+00 |
| DFN8 | $\bar{r}(nm)$ | N/A | N/A | N/A | 6.950e+00 | 2.644e-03 | 7.212e-04 |
| DFN8 | $V_p(nm^3)$ | 6.704e+02 | N/A | 1.905e+00 | 7.020e+02 | 1.303e+01 | 3.752e+00 |
| DFN8 | $V_c(nm^2)$ | 1.714e+01 | N/A | 8.717e-03 | 1.709e+01 | 1.678e-01 | 2.065e-02 |
| DFN8 | $\ell_c(nm)$ | N/A | N/A | N/A | 6.540e+00 | 1.374e-01 | 2.735e-02 |
| DFQ8 | $R_g(nm)$ | 2.745e+00 | 1.442e-03 | 4.122e-03 | 2.748e+00 | 4.199e-03 | 1.135e-03 |
| DFQ8 | $I(0)$ | 1.542e+04 | 5.665e+00 | 1.017e+01 | 1.542e+04 | 4.570e+00 | 4.596e+00 |
| DFQ8 | $\bar{r}(nm)$ | N/A | N/A | N/A | 3.594e+00 | 3.010e-03 | 1.164e-03 |
| DFQ8 | $V_p(nm^3)$ | 9.769e+01 | N/A | 3.367e-01 | 9.917e+01 | 1.278e+00 | 4.373e-01 |
| DFQ8 | $V_c(nm^2)$ | 4.638e+00 | N/A | 2.890e-03 | 4.355e+00 | 2.949e-02 | 4.579e-03 |
| DFQ8 | $\ell_c(nm)$ | N/A | N/A | N/A | 3.624e+00 | 5.274e-02 | 1.227e-02 |

It is important to note that the statistical errors described here are only based on resampling the scattering profile and do not account for systematic error that is likely to dominate. As discussed above, *V_p_, V_c_*, and *ℓ_c_* are particularly sensitive to systematic deviation. To test the algorithm for accuracy with experimental data, we calculated *V_p_* values for 29 datasets from the Benchmark section of the SASBDB and used *V_p_* to estimate the molecular weight (MW) of the particle (where *MW* = VP/1.6). Figure 7 shows the comparison of molecular weight values calculated using *V_p_* estimates from *denss.fit_data.py* and DATPOROD and compared with their expected values. Here, the expected value is taken from the expected molecular weight in the SASBDB entries calculated from the amino acid sequence. The median error from *denss.fit_data.py* is 8.7% and from DATPOROD is 18.0%. As expected, these real errors in practice are significantly larger than the < 2% statistical or estimated errors in Table 2, confirming that systematic deviations dominate actual estimates of Porod volume from experimental data.

**Fig. 7.**
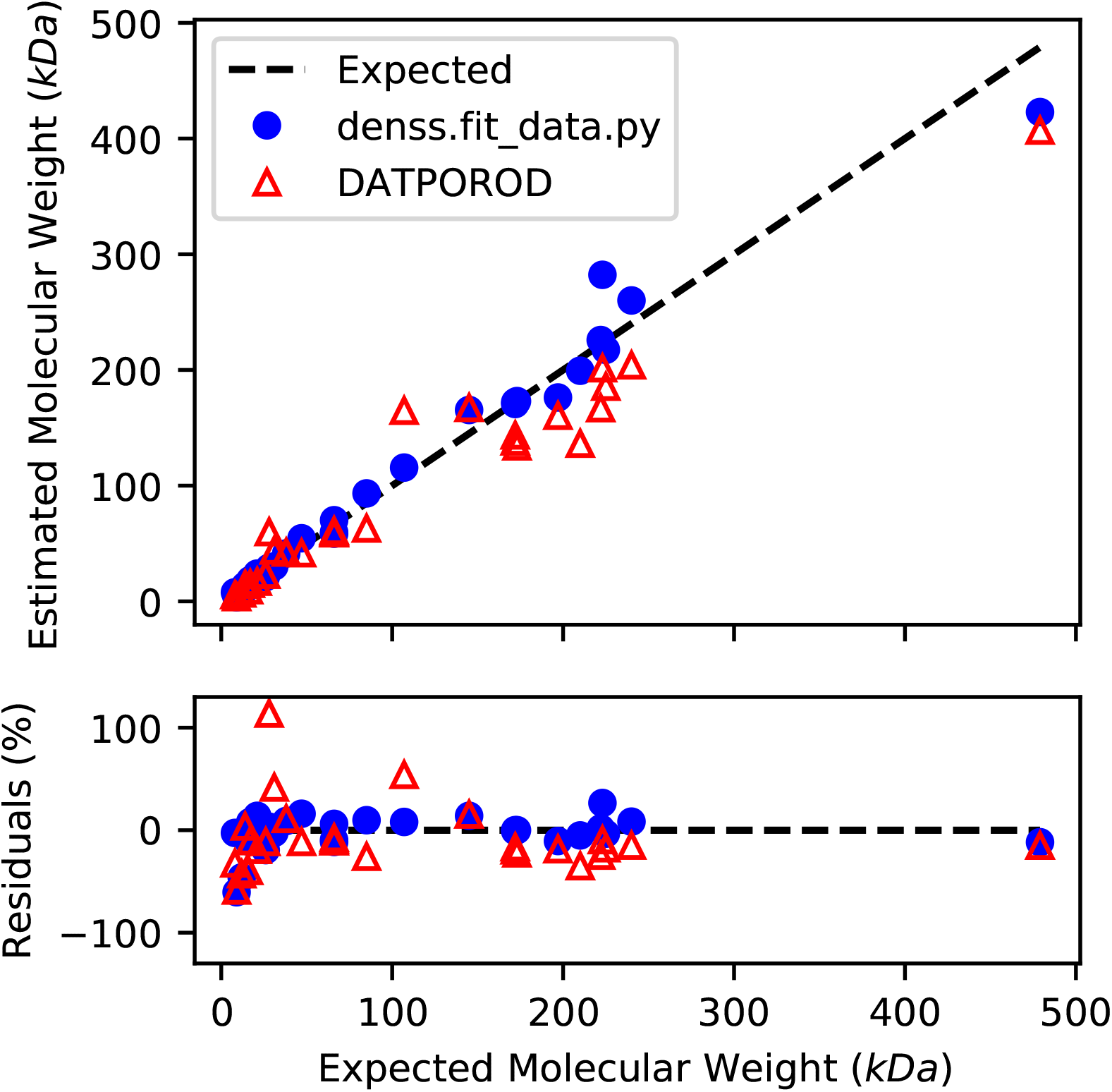
Comparison of expected molecular weight with values calculated using *V_p_* estimated from *denss.fit_data.py* and DATPOROD.

## 4. Discussion

The approach outlined above is an extension of Moore’s original description of SAS profiles using a trigonometric series with the advantage of replacing the nondescript Moore coefficients with specific intensity values. As such, this derivation is subject to all of the same requirements as Moore’s, including the need for accurate intensity measurements for at least the first three Shannon channels to obtain reliable estimates of parameter values. We have described a derivation for performing regularization of the real space *P*(*r*) curve analytically, and procedures for the automatic estimation of *D* and *α* values. We also present a novel approach for estimating parameters that are particularly sensitive to systematic deviations at high *q* values, such as *V_p_*.

As in Moore’s original approach, the use of least squares minimization for the derivation above of a series of SAS parameters directly from the *I_n_* values enabled the estimation of uncertainties through error propagation while accounting for covariances in the data. The oversampling of the information content in the SAS profile effectively increases the signal to noise ratio of each of the unique observations in the data, i.e. the *I_n_*’s. Additionally, the analytical regularization derived above simultaneously enables smooth *P*(*r*) curves and accurate fits to experimental data, all while providing error estimates for the *I_n_*’s and associated parameter calculations accounting for covariances in the data. Using simulated and experimental data, we show that these methods yield parameter values describing the size and shape of particles that are as accurate, or often more accurate, than existing tools.

The algorithm has been made available open-source as a script called *denss.fit_data.py*, accessible on GitHub at https://github.com/tdgrant1/denss. The software can be run either from the command line or as an interactive GUI.

## Appendix A Extension of Moore’s IFT

Moore uses a trigonometric series to define a function *Q*(*r*) = *P*(*r*)/*r*. This definition resulted in a convenient relationship between the real space *Q*(*r*) and the reciprocal space *U*(*q*) = *q_I_*(*q*), where the two are Fourier mates. This results in equations 17 through 18 defining *P*(*r*) and *I*(*q*):

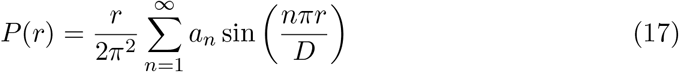

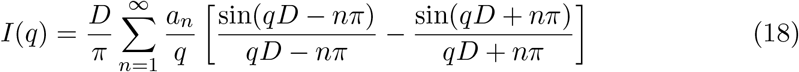

where *a_n_* are weights for each term in the series, the Moore coefficients, and *D* is the maximum particle dimension (Note: modest variations compared to Moore *s* original description of these functions by a factor of 2*π* are due the use of *q* = 4*π* sin(*θ*)/*λ* rather than *s* = 2 sin(*θ*)/*λ*, where 2*θ* is the scattering angle and *λ* is the wavelength). Key to Moore s approach (and other IFT methods (Glatter, 1977; Svergun, 1992)) is that the weights *a_n_* define both the real space and reciprocal space profiles, using the appropriate basis functions. Least squares can be used to determine the *a_n_*’s and the associated standard errors by minimizing the *χ*^2^ formula (equation 19):

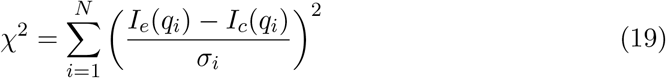

where *I_e_* is the experimental intensity for data point *i, I_c_* is the intensity calculated at *q_i_* given by equation 18, *σ_i_* is the experimental error on the intensities, and *N* is the total number of data points.

Moore’s use of Shannon information theory to define *I*(*q*) resulted in a selection of *q* values, namely *q_n_* = *nπ/D*, termed “Shannon channels” (Feigin & Svergun, 1987; Svergun & Koch, 2003; Rambo & Tainer, 2013). The intensities at *q_n_*, i.e. *I_n_* = *I*(*q_n_*), therefore become important values as they determine the *a_n_*’s and thus can be used to completely describe the low-resolution size and shape of a particle obtainable by SAS. It is therefore convenient to derive the mathematical relationship between In and *a_n_*. Note that here we will further use *m* to refer to a particular term in the series, and we will use *n* when referring to the terms in the function defining the entire series. The intensity *I_m_* at *q_m_* = *mπ/D* is

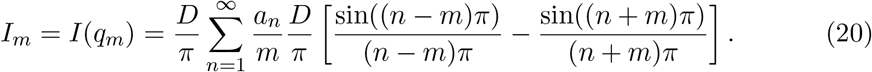

Since

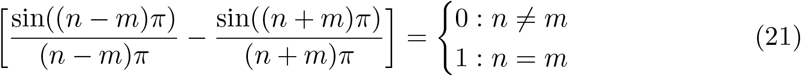

the sum reduces to a single term when *m = n*, resulting in

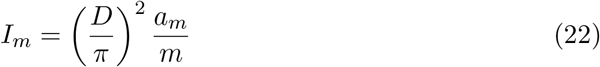

and therefore

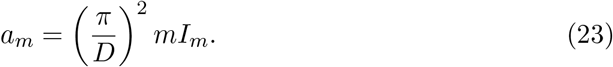

Equation 23 defines a relationship between the mth Moore coefficient and the intensity at the *m*th Shannon point. Inserting equation 23 into equation 18 and simplifying yields a general equation for *I*(*q*) as a function of the intensity values at the Shannon points:

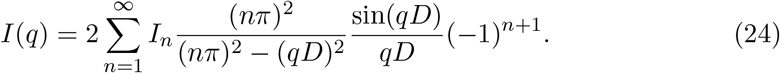

Defining the basis functions *B_n_* as

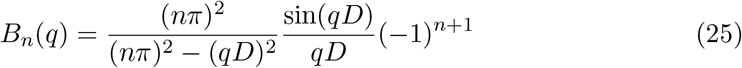

*I*(*q*) can now be expressed as a sum of the basis functions *B_n_* weighted by physical intensity values at *q_n_*

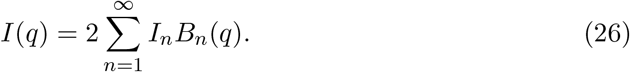

As in Moore’s original approach, the *B_n_* functions are determined by the maximum dimension of the particle, *D. B_n_*’s for *D* = 50 Å are illustrated in Figure 1. The *P*(*r*) function can be determined from the continuous *I*(*q*) according to equation 27:

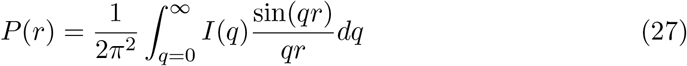

*P*(*r*) can also be represented using the series of *I_n_* values by inserting equation 23 into equation 17, resulting in equation 28:

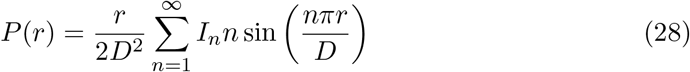

or by defining real space basis functions *S_n_* as follows:

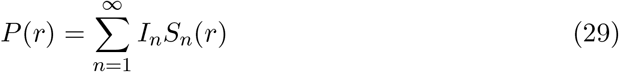

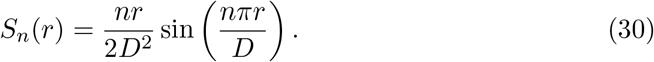

As intensity values measured precisely at each *q_m_* are typically not collected during experiment, least squares minimization of *χ*^2^ can be used to determine optimal values for each *I_m_* from the oversampled SAS profile, resulting in greatly increased precision for each *I_m_* compared to measured intensities. To determine the set of optimal *I_m_* values, let

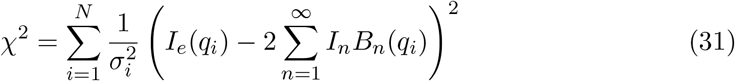

Values for each *I_m_* are sought which minimize *χ*^2^, i.e. where *δχ*^2^/*δI_m_* = 0, yielding

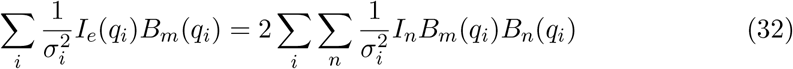

for all *m*. Let

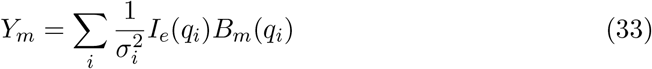

and

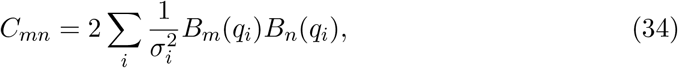

then ***Y*** = ***IC*** and ***I*** = ***YC***^-1^, where ***I*** and ***Y*** denote the arrays of *I_m_* and *Y_m_* values, respectively, and **C** is a matrix whose elements are *C_mn_*. The *I_m_*’s can now be used with equation 26 to provide a smoothed representation of *I*(*q*) as well as *P*(*r*) using equation 29. The matrix **C**^-1^ contains all the information on the variances and covariances of **I**. The standard deviation for *I_m_* can thus be calculated from **C** as

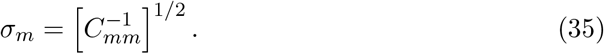

Furthermore the errors on the calculated *I_c_*(*q*) curve can be calculated as:

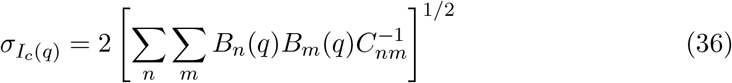

and the errors in *P*(*r*) are

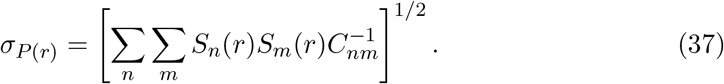

## Appendix B Derivation of Size Parameters and Error Estimates

Here the detailed derivation is presented for calculating *R_g_* from the *I_n_*’s. Derivations for the remaining parameters and errors can be determined similarly. *R_g_* can be calculated from the *P*(*r*) curve according to the following equation:

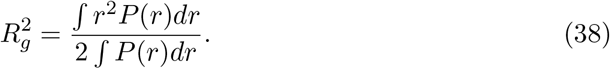

To determine the equation relating *I_n_* coefficients to *R_g_*, we substitute 28 into equation 38. Since

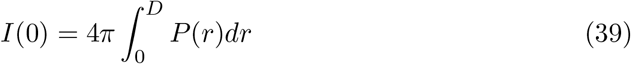

the denominator can be simplified to

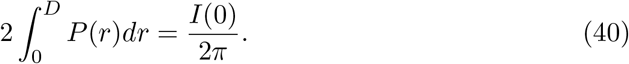

The numerator becomes

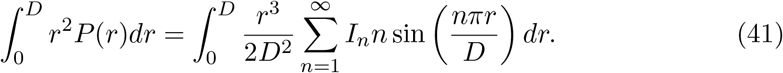

Since Porod’s law shows that intensity (and thus the *I_n_*’ s) decays as *q*^-4^ for globular particles, and similarly as *q*^-2^ for random chains, the infinite sum is guaranteed to converge to a finite value. Since the sum converges, the Fubini and Tonelli theorems (Fubini, 1907; Tonelli, 1909) show that the infinite sum can be exchanged with the finite integral as follows

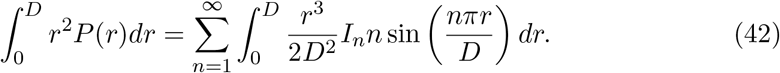

Pulling constants out of the integral results in

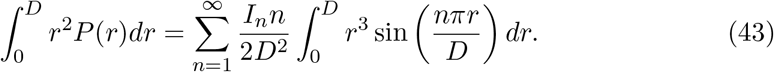

The integral can be solved by three iterations of integration by parts and evaluated at the limits to obtain

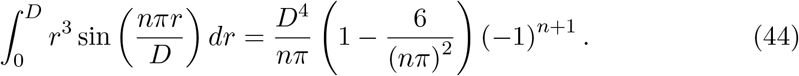

Equation 44 can be combined with equation 43 and simplified to obtain the following equation for the numerator

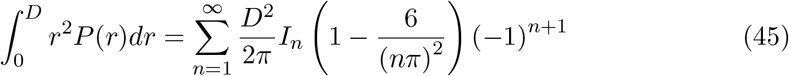

which can be combined with equation 40 to ultimately obtain equation 8. Similar steps can be followed for the remaining parameters.

The average vector length in the particle, 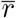, is defined as

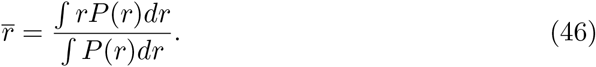

which can be combined with equation 28 to yield equation 10.

The Porod invariant, *Q*, is defined as the integrated area under the Kratky plot (Porod, 1982)

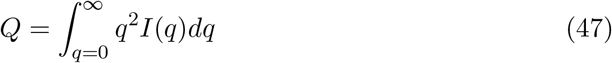

which, by plugging in equation 26, yields equation 12.

The Porod volume can then be calculated, using its definition containing the Porod invariant (Porod, 1982), by the following equation

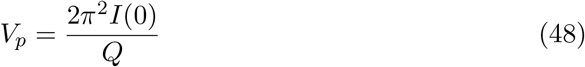

The Volume of Correlation (Rambo & Tainer, 2013), *V_c_*, is defined as

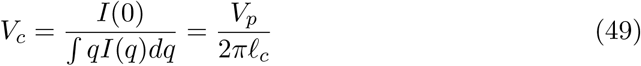

where *ℓ_c_* is the length of correlation (Porod, 1982). *V_c_* can thus be estimated from the *I_n_*’s according to equation 13. The length of correlation can be found from equation 49 or by combining equations 47, 48 and 49 it can be shown that

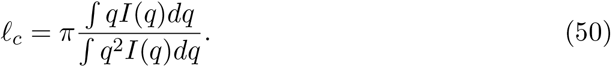

Plugging in equation 26 into equation 50 yields equation 14.

Since the matrix ***C***^-1^ contains all the information on the variances and covariances of the *I_n_*’s, the uncertainties in each parameter can be derived using error propagation. The error in *I*(0) is thus

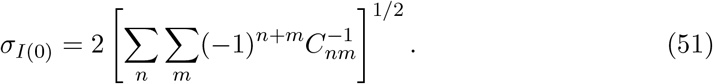

The error in *R_g_* is

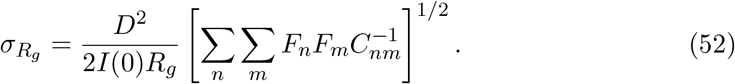

In equation 8 it can be seen that *R_g_* has a non-linear dependence on *I_n_*, and thus the error on *R_g_* is dependent on *R_g_* itself. The error in 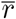 is

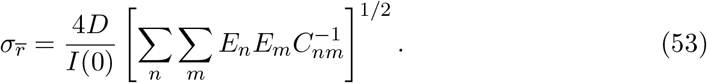

The error in *Q* is

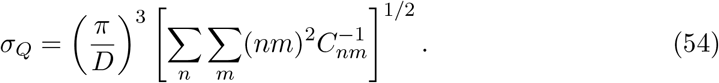

The error in *V_p_* is

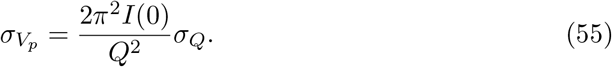

The error in *V_c_* is

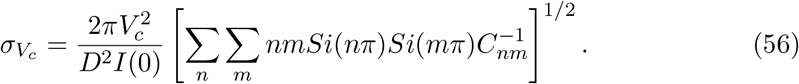

The error in *ℓ_c_* estimated from equation 49 becomes

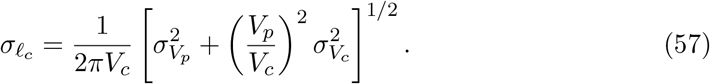

## Appendix C Derivation of Analytical Regularization of *P*(*r*)

To enable the regularization of the *P*(*r*) curve for the derivation described above, *S* has been chosen to take the commonly used form of equation 58:

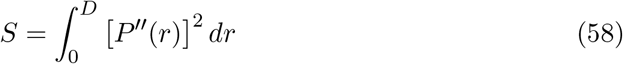

where *P*”(*r*) is the second derivative of *P*(*r*) with respect to r. The second derivative is often chosen as it is sensitive to large oscillations in the *P*(r) function, i.e. smoother functions will have fewer oscillations and thus *S* will be small. This representation allows for an analytical solution to the problem of regularization of the *P*(*r*) curve. To begin, the second derivative of *P*(*r*) can be calculated as

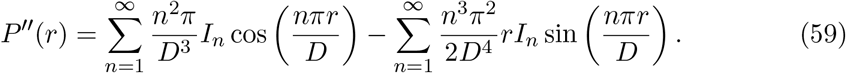

Equation 58 requires squaring and then integrating equation 59, which is not possible analytically. However, ultimately we want to use least squares minimization of *T* in equation 15 to determine the optimal *I_m_* values that yield the best fit of *I*(*q*) to the experimental data and a smooth *P*(*r*) function. The minimization of *T* with respect to *I_m_* requires taking the derivative of S with respect to *I_m_*. Therefore, rather than attempting to square and integrate equation 59, we can first take the derivative of S with respect to *I_m_* as follows:

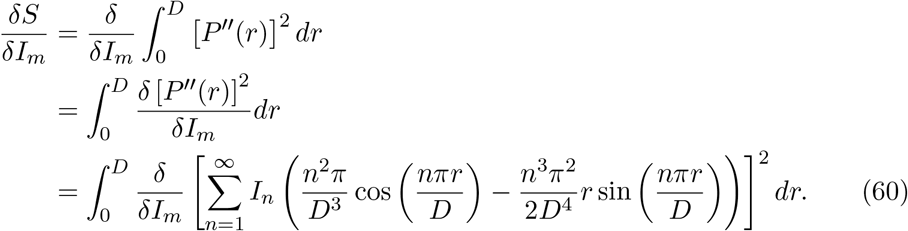

Now we can take the derivative as

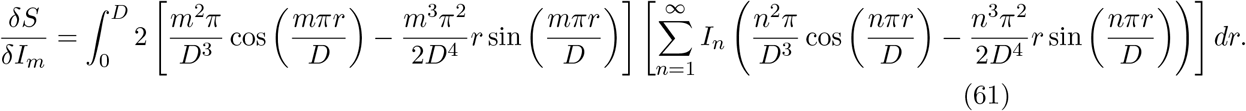

Since the term in square brackets outside the sum is independent of *n*, it can be brought inside the sum, and the integration and summation exchanged:

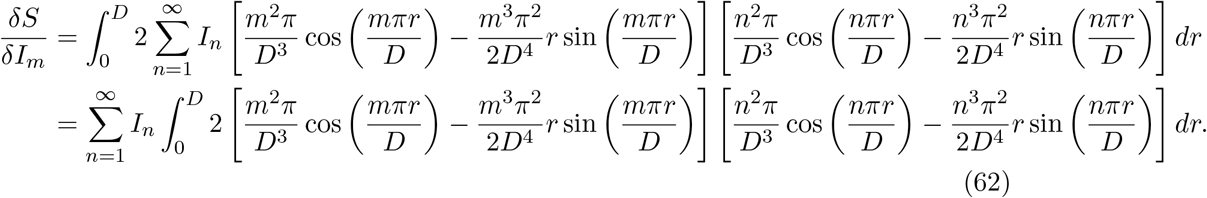

The terms in the integrand can now be expanded, yielding four terms in total:

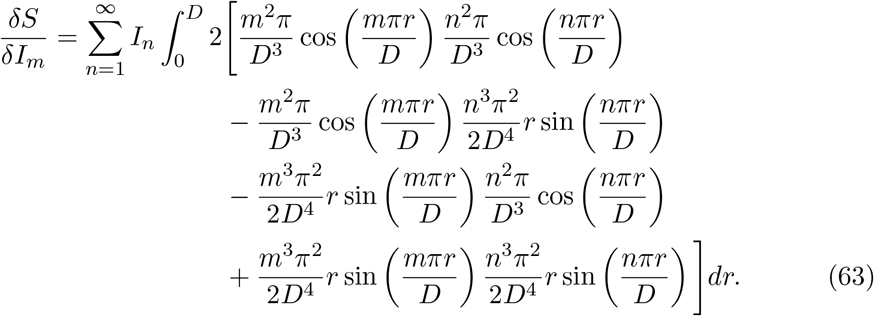

Each term can now be integrated and evaluated at the limits to obtain the following:

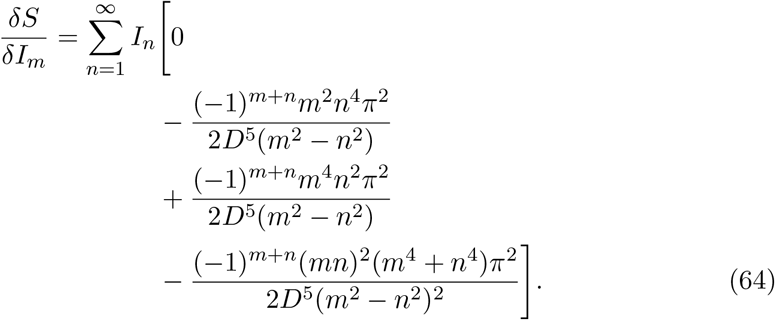

These terms can be combined and represented by *G_mn_* below. Note that when *m* = *n* the equation is undefined, so the integration has been repeated after taking the limits as *m* approaches *n* for the special case when *m = n*. Taken together, the derivative of *S* with respect to *I_m_* can be now be represented as equation 65:

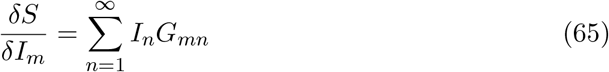

where

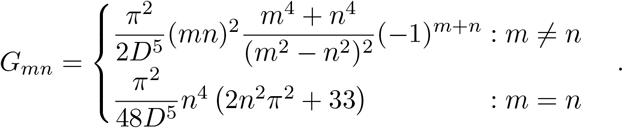

Following the same procedure outlined in equations 31 through 34 and now including 65, equation 15 can now be solved by least squares minimization to yield the optimal values for each *I_m_* while accounting for the regularizing function S according to the following modified equations:

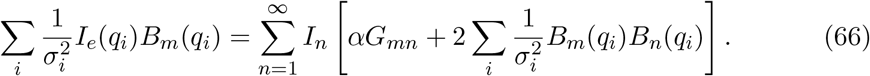

Once again, letting

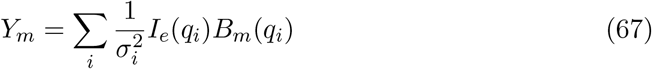

and now letting

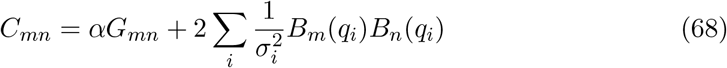

then the matrix of *I_m_* values, ***I***, can be found in a similar fashion as before:

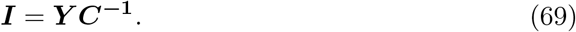

## Appendix D Derivation of Sphere Parameters

The equation governing the scattering of a sphere of radius R is given by equation 70 (Rayleigh, 1911; Porod, 1982):

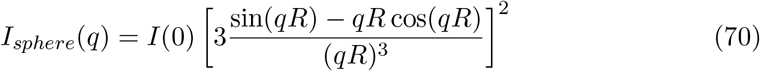

(for simplicity, *I*(0) is here used as a global scale factor accounting for proportionality constants related to volume, concentration, scattering length density, etc., and is set to unity in the following equations). By evaluating equation 70 at *q_n_* = *nπ/D* = *nπ*/2*R*, we obtain a representation of the intensity from a sphere as a function of *I_n_*’s, shown by equation 16. All of the parameters derived above can be expressed for a sphere as well, yielding well-known values for spheres of radius *R* = *D*/2. For example, equation 8 can be used to calculate the radius of gyration of a sphere of radius *R* by inserting equation 16 yielding

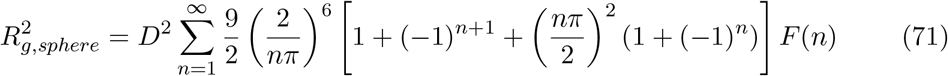

The entire infinite sum converges to 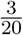 and the expression simplifies to

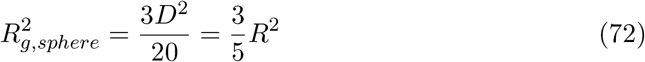

which is the well-known equation relating the radius of gyration of a solid sphere to its radius. Similarly, the Porod invariant *Q* of a sphere can be calculated according to equation 12 and shown to be

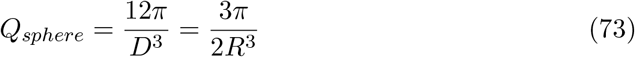

which can be combined with equation 48 to evaluate the Porod volume as

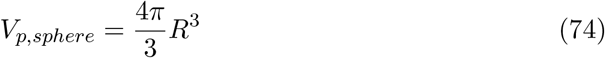

which is the well-known equation for the volume of a sphere of radius *R*. Each of the other parameters listed in the equations above can also be solved in a similar fashion and shown to be as follows:

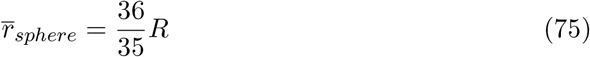

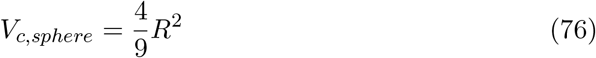

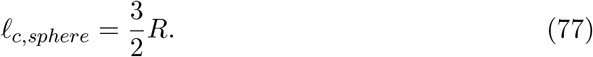

The *P*(*r*) curve for a sphere has previously been derived (Porod, 1982):

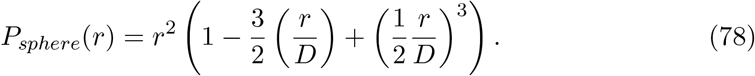

## Acknowledgements

The author thanks Drs. Stephen Meisburger, Kushol Gupta, and Robert Rambo for testing the software and for useful discussions.

## Synopsis

An indirect Fourier transform method is presented which describes a solution scattering profile from a reduced set of intensities. Equations are derived to fit the experimental profile using least squares and to calculate commonly used size and shape parameters directly from the reduced set of intensities, along with associated uncertainties. An analytical equation is derived enabling regularization of the real space pair distribution function. Convenient software is provided to perform all described calculations.

